# Delta-range coupling between prefrontal cortex and hippocampus supported by respiratory rhythmic input from the olfactory bulb in freely behaving rats

**DOI:** 10.1101/2020.05.04.077461

**Authors:** Rola Mofleh, Bernat Kocsis

## Abstract

An explosion of recent findings firmly demonstrated that brain activity and cognitive function in rodents and humans are modulated synchronously with nasal respiration. Rhythmic respiratory (RR) coupling of wide-spread forebrain activity was confirmed using advanced techniques, including current source density analysis, single unit firing, and phase modulation of local gamma activity, creating solid premise for investigating how higher networks use this mechanism in their communication. Here we show essential differences in the way prefrontal cortex (PFC) and hippocampus (HC) process the RR signal from the olfactory bulb (OB) allowing dynamic PFC-HC coupling utilizing this input. We used inter-regional coherences and their correlations in rats, breathing at low rate (∼2 Hz) at rest, outside of the short sniffing bouts. We found strong and stable OB-PFC coherence, contrasting OB-HC coherence which was low but highly variable. PFC-HC coupling, however, primarily correlated with the latter, indicating that HC access to the PFC output is dynamically regulated by the responsiveness of HC to the common rhythmic drive. This pattern was present in both theta and non-theta states of waking, whereas PFC-HC communication appeared protected from RR synchronization in sleep states. The findings help to understand the mechanism of rhythmic modulation of non-olfactory cognitive processes by the on-going regular respiration, reported in rodents as well as humans. These mechanisms may be impaired when nasal breathing is limited or in OB-pathology, including malfunctions of the OB epithelium due to infections, such as in COVID-19.

## 1. Introduction

More than half a century after the first observations ^1^, an explosion of findings firmly demonstrated that brain activity and cognitive function in rodents and humans are modulated synchronously with nasal respiration (rev. ^2^). Respiratory related oscillations (RRO) were detected in numerous brain structures, including higher order cognitive centers in the prefrontal cortex (PFC)^3, 4^ and hippocampus (HC)^5-7^. Respiratory rhythm primarily derives from rhythmic nasal airflow of the olfactory bulb (OB)^6^, which dynamically couples with intrinsic network oscillations in these structures either: (1) by coherence, when the frequency of RRO matches that of local field potentials (LFP) such as delta and theta activity in rodents ^3, 5-9^, or (2) by phase-amplitude modulation when the frequencies diverge, as in gamma oscillations in rodents ^3, 4, 6, 8^. In human, where respiratory rate (< 0.5 Hz) is out of the frequency range of most EEG rhythms, relevant for cognitive function (delta, theta, alpha and gamma oscillations), RRO is also established using the mechanism of phase-amplitude modulation^10^.

Accumulating evidence over the past decade has advanced research on the mechanisms underlying OB-cortical RRO coupling from well-studied sniffing episodes, to mechanisms associated with continuous on-going respiration – thereby raising questions about how RRO may be involved in non-olfactory cognitive processing ^11, 12^. Respiratory modulation of a wide range of cognitive functions has been reported both in rodents and human, from sensory processing and motor coordination to various memory functions (rev. ^12^) – which are not directly related to olfaction or to gas exchange (as a primary respiratory function). Rhythmic coupling is a powerful, ubiquitous mechanism of functional coordination of neural ensembles, and RRO appears to be a potential source of wide, perhaps even global ^2, 9^ synchronization of various networks, cortical as well as subcortical. For functional networks, access to rhythmic drive has to be dynamically regulated in a state- and task-dependent manner to encompass specific circuits involved in particular tasks – to couple them when necessary and uncouple them when it is not. The anatomical systems carrying the RRO signal to diverse networks appear suitable to exert such control. RRO from different sources ^13^, of which OB is dominant, is transmitted to various networks which may differentially synchronize with this input dependent on their unique circuit characteristics and connectivity. For example, key intrinsic oscillations in PFC (delta, 2-5 Hz ^14-16^) are in the range of on-going RRO, whereas those in the HC are faster (theta, 5-10 Hz ^17^), overlapping in rodents with sniffing frequency. These two forebrain structures are the focus of the current study investigating how the RRO signal generated in the OB may potentially contribute to PFC-HC communication by synchronizing their activities at the respiratory rate.

Effective PFC-HC communication is important for normal cognition and impaired PFC-HC coupling was implicated in cognitive deficits in psychiatric diseases ^18-20^. We have shown recently that rhythmic coupling between PFC and HC can be established in both delta and theta ranges, or even simultaneously, and proposed that they may serve as parallel channels of communication (but in opposite directions) between the two structures ^21^. RRO and theta was shown to co-exist during theta states with an asymmetric regional distribution in the cortex; that is, RRO dominant in the frontal cortex and theta in more caudal cortical areas ^9^. Based on these data we hypothesized that RRO might enhance the communication between PFC and HC networks primarily in the PFC-to-HC direction. Indeed, we found a strong and reliable RRO transmission through the OB to PFC whereas OB-HC coherence was low in all states. RRO in HC was highly variable but showed strong correlation with RRO coherence between PFC and HC. Thus, PFC delta output was steadily segmented and shaped by the RRO, but it was the HC response to the common RRO drive that dynamically regulated PFC-HC coupling in the delta range. The details of this regulation remain unknown. Importantly however, the strong effect of RRO on cortical coupling suggests that damage to OB may lead to functional abnormalities in higher brain function. It is known for example that SARS-CoV-2 viral infection of ACE2 receptor-expressing epithelial cells in the OB leads to loss of smell ^22^, associated with significant MRI and correlated with not only smell but also with memory loss ^23^. A similar mechanism may affect the OB-dependent RRO as well.

## 2. Methods

Male rats (360–560g, Charles River Laboratories) were used in this study. Experiments were performed on 8 rats subjected to survival surgery followed by chronic recordings in free behaviors. All procedures were approved by the Institutional Animal Care and Use Committee of Beth Israel Deaconess Medical Center and carried out in accordance with relevant guidelines and regulations. The study and reporting also adheres to ARRIVE guidelines.

### 2.1. Experimental procedures

Diaphragmal EMG was recorded in all rats along with LFP in the OB, PFC and HC using microwires (Fig. S5) and electrocorticograms using screw electrodes over the parietal cortex with additional EMG recordings in the neck muscle (see Supplementary Information for details). Recordings were made in undisturbed condition. Recordings started 7-10 days after surgery in two 24 hr recording sessions, acquired a couple of days apart in each rat. Sleep-wake states were identified using standard criteria based on cortical EEG, HC LFP, and neck muscle EMG recordings. For analysis, multiple segments were selected from discontinuous episodes of each state dispersed over the 2 days of recordings, in which respiration appeared relatively stable without fluctuations (Table S1). Respiratory rate varied in different states in a relatively narrow band (between 1-3 Hz, Table S1) with an occasional faster component (4-6 Hz) in AW which did not overlap with theta frequency, specifically verified in each segment submitted for analysis.

### 2.2. Data analysis

was performed on recordings acquired at 1 kHz. Dia EMG recordings were processed using built-in procedures of Spike2 to remove ECG contamination and to convert high-frequency EMG components in order to retrieve pure respiratory rhythm. Noise-free segments with stable respiration for at least ∼100 s were selected in slow wave sleep (SWS), rapid eye movement sleep (REM), quiet waking (QW), and active waking (AW), recorded on two different days (see Table S1 for the number and length of segments in each state), and submitted to Fast Fourier Transform to obtain power spectra and coherence function with ∼0.25 Hz frequency resolution. Coherence values were compared against chance using surrogate-based statistical testing ^9^ and potential contamination due to volume conduction or common reference was tested using cross-correlations in the time domain (see more details in Supplement, Fig.S6). To quantify neuronal synchronization between different structures we used pairwise coherences, calculated between 4 signal pairs, representing the potential transfer of the RRO signal to higher-order structures through the OB (i.e. dia with OB and OB with PFC and HC) and between these higher order structures (i.e. PFC with HC). Power spectra for dia EMG and HC were also calculated to identify the frequencies of spectral peaks of RRO and theta rhythm. Coherence values at RRO and theta frequencies were calculated in each segment (Table S1) and daily averages of these values were used in statistical analysis, including group averages (Tables 1 and S2) and comparisons of peak coherences. Differences between coherences in different states were tested using Student’s t-test after Fisher r to z transformation to obtain z-scored values with normal distribution. Correlation between pair-wise coherences was statistically tested using Excel’s T-DIST procedure (see more details in Supplement).

**Table 1.** Relationship between coherences through OB and their effect on RRO synchronization between PFC and HC. Yellow background highlights relatively high coherences, above 0.4 and significant correlations (p < 0.05), R^2^, between pairwise coherences. C(X-Y): coherence between X and Y.

| Peak Coherence | SWS | REM | QW | AW |
| --- | --- | --- | --- | --- |
| C(Dia-OB) | 0.29±0.04 | 0.33±0.04 | 0.66±0.05 | 0.40±0.07 |
| C(OB-PFC) | 0.25±0.04 | 0.32±0.04 | 0.67±0.05 | 0.55±0.07 |
| C(OB-HC) | 0.18±0.05 | 0.09±0.03 | 0.29±0.07 | 0.24±0.10 |

**B.** Correlations between C(HC-PFC) and coherences connecting OB to other signals
| $R^2$ between C(PFC-HC) and... | SWS | REM | QW | AW |
| --- | --- | --- | --- | --- |
| ... C(Dia-OB) | 0.08 | 0.01 | 0.00 | 0.00 |
| ... C(OB-PFC) | 0.04 | 0.07 | 0.33 | 0.12 |
| ... C(OB-HC) | 0.27 | 0.00 | 0.69 | 0.79 |

**C.** Coherences connecting LFP signals of PFC and HC, at RRO frequency
| peak coherence | SWS | REM | QW | AW |
| --- | --- | --- | --- | --- |
| C(PFC-HC) | 0.22±0.04 | 0.19±0.07 | 0.45±0.08 | 0.42±0.09 |

## 3. Results

### 3.1. Respiratory rhythm in diaphragmic EMG correlates with LFP oscillations in OB

Respiratory rate varies extensively in rodents, covering the entire range of characteristic frequencies of low frequency oscillations (delta, theta, even alpha) intrinsically generated by neural circuits in the cortex and hippocampus ^2^. To focus on on-going regular RROs (Fig. 1A), the present analysis was limited to lasting stationary segments – excluding short segments with fast RROs potentially associated with sniffing (Fig. S1). Thus, RRO frequency was in the delta range in all states – below 2Hz in sleep, and slightly faster in waking, but still below the theta range (Table S1). Except for (AW), RRO was stable in all states, manifested by a single sharp peak in the autospectra of the dia EMG signal in each recording session (Fig. S2A). RRO frequency in these states (QW, REMs, SWS), shifted from experiment to experiment in a narrow range, producing ∼1 Hz-wide peaks in average dia autospectra (Fig. 1B). LFP in the OB was correlated with this signal in a state-dependent manner (see below) giving rise to RRO peaks in the dia-OB coherence spectra, which in sleep were also restricted to this narrow frequency range (Fig 1C).

**Figure 1.**
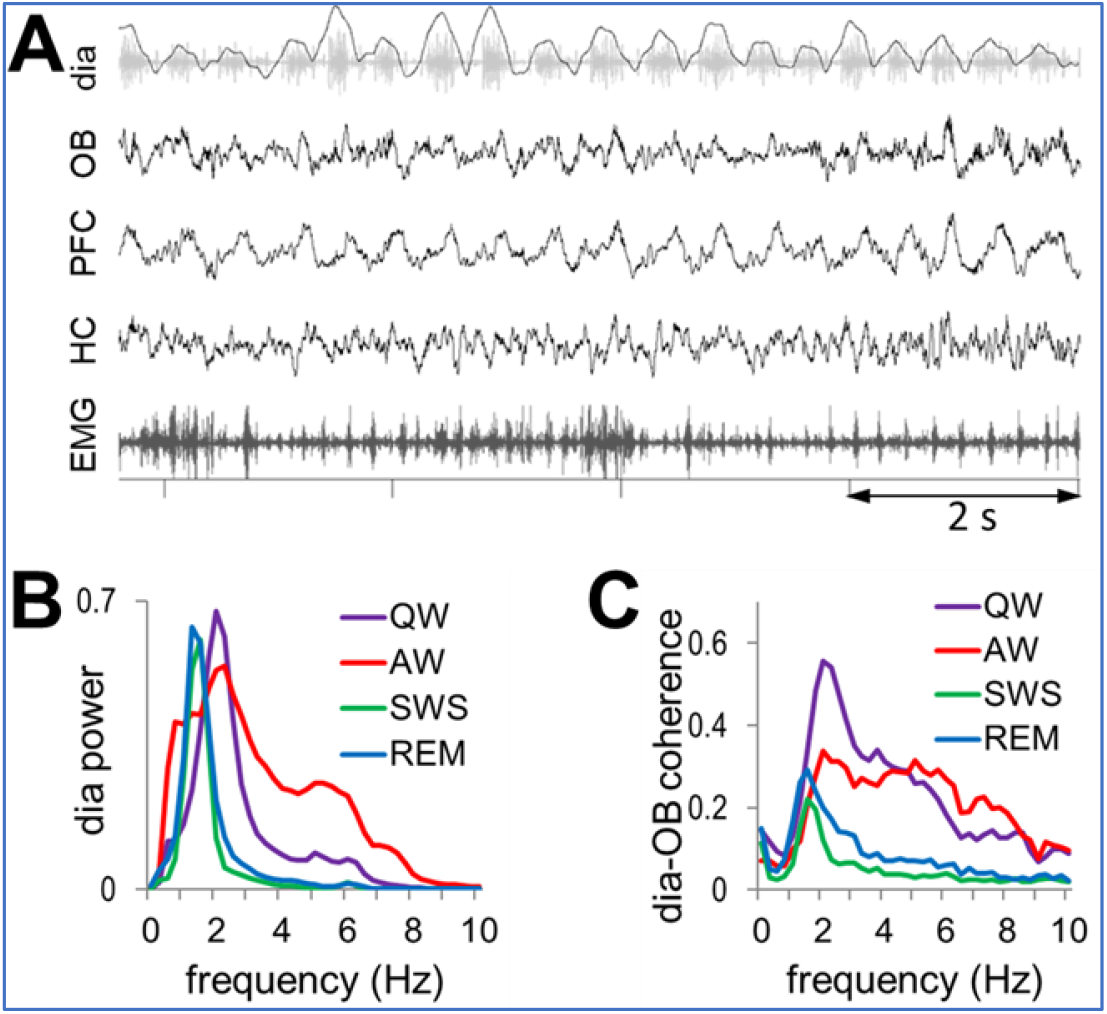
**A**. Sample recording of respiratory rhythm (black) derived from diaphragmal EMG (grey) along with LFPs in OB, PFC, and HC and neck muscle EMG in QW state. **B**. Group averages of dia autospectra in different states. Note narrow RRO peaks in all recordings at ∼2 Hz. Power is shown in arbitrary units after normalization of autospectra in individual recordings setting maxima equal to 1. **C**. Group averages of dia-OB coherence spectra, in different states. Note coherence peaks constrained to RRO frequencies (dia spectral peaks) in sleep and in a wider range, up to 6 Hz in wake states. In AW, dia-OB coherence does not have a clear RRO peak on the group average due to interindividual variability of the respiratory rates (see in Fig. S2).

In waking, RRO peaks in dia autospectra were somewhat wider in each experiment (indicating short-time scale variations within recording sessions) and its peak shifted in a wider range between experiments (1-3 Hz; Fig. 1B). In a few recordings (4 in AW and 1 in QW), there was a second dia power peak at 4-6 Hz (Fig. S2A), but this always appeared together with a clearly distinguishable 1-3 Hz component (Fig. S1C). The lower peak (1-3 Hz) was also present in the group averages of dia power spectra (Fig.1B) and dia-OB coherence functions (Figs.1C, S2B).

To examine the origin of RRO coupling between higher order networks, we used pairwise coherences between dia, OB, PFC, and HC signals and their correlations, calculated at the respiratory frequency in all states. Respiratory rate was identified from dia autospectra, in each individual experiment.

### 3.2. OB unevenly conveys RRO to higher order networks in PFC and HC

To assess RRO synchronization across regions, RRO peaks of coherences between signal pairs representing RRO transfer from rhythmic nasal airflow to the OB and then from OB to PFC and HC, were compared in different behavioral states. dia-OB coherences showed strong state dependence (Fig. 1C and 2A, Table 1A) and while RRO coherence between OB and PFC followed this pattern, those between OB and HC were relatively low in all states (Figs. 2A). Thus, OB-PFC coherences at RRO frequency were higher during wake than sleep states; differences between AW and QW vs. SWS and REMs were all statistically significant (p<0.01), whereas within waking and within sleep no significant differences were detected (p>0.1). On the other hand, group averages of OB-HC coherence were in a narrow range; they did not change between QW, AW, and SWS and were somewhat lower in REM sleep (p<0.1). In all states, OB-PFC coherences were similar to dia-OB coherence (i.e. statistically equal, p>0.1), in major contrast to the pathway conveying RRO to HC; where OB-HC and dia-OB coherences were significantly different in all states (p<0.01), except SWS.

**Figure 2.**
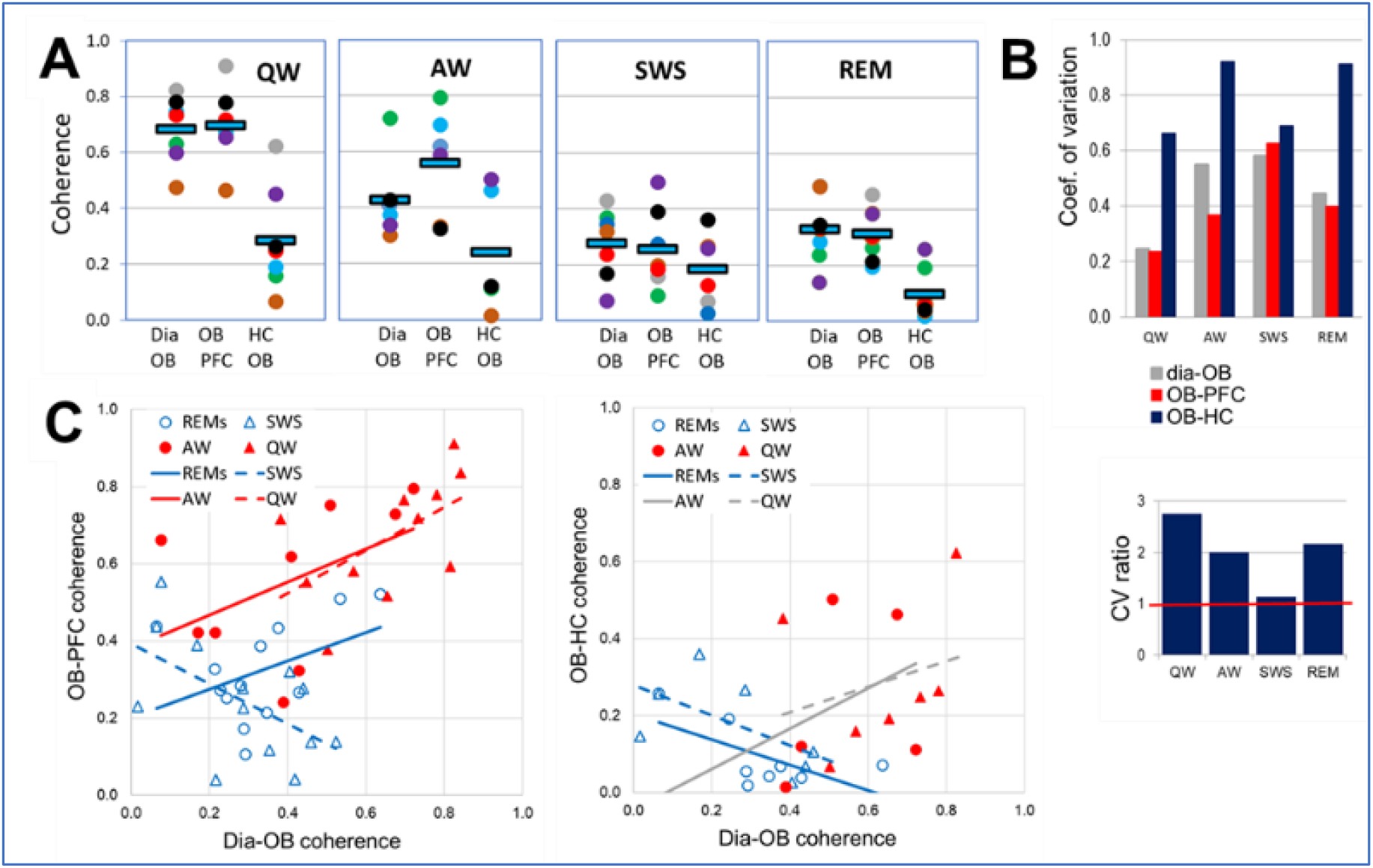
Comparison of state-dependent RRO coherences in PFC and HC transferred through OB. **A**. Peak coherence at RRO frequency between rhythmic dia activity and LFP in the OB and between OB and cortical (PFC) and hippocampal (HC) networks during different sleep-wake states. Note strong state dependence and nearly identical dia-OB and OB-PFC coherences in all recordings and considerably lower OB-HC coherence. Squares: group averages, dots: individual experiments; same colors identify individual rats. **B**. Variability of coherence values in individual experiments in different states. Coefficient of variation (top) and CV ratio (bottom) of coherences in the OB-HC vs. the other two signal-pairs (dia-OB and OB-PFC). Note high variation of OB-HC in waking (AW and QW) and REM sleep, 2-3 times exceeding CV of the other pairs. **C**. Relationship between peak RRO coherences connecting dia to OB (Dia-OB) and those connecting OB to neural networks of PFC (left) and HC (right) in different states. Trend-lines with non-significant correlations (p>0.1) are shown in grey; solid lines show theta, dashed line show non-theta states. Note significant positive correlation of between dia-OB and OB-PFC, but no positive correlation between dia-OB and OB-HC coherences.

Examination of pairwise coherences in individual experiments further supported the pattern revealed by group averages; OB-HC coherence was lower than dia-OB and OB-PFC in all experiments in all states (see colors of dots, representing different experiments in Fig. 2A), even though the increase in RRO coherences during wake states compared to sleep, showed natural variation between experiments. Furthermore, the variability of coherence values revealed a feature, unique for the OB-HC relationship, further separating it from dia-OB and OB-PFC. In QW and AW, the coefficient of variation (CV; Fig. 2B) reflected widely dispersed values of OB-HC coherences, compared with the other signal-pairs (its CV exceeded that of dia-OB and OB-PFC by 100-200 %; CV ratio>2; Fig. 2B).

To study the origin of this variability, dia-OB coherences quantifying RRO input to the OB were correlated with RRO coherences in the pathways connecting OB further to the PFC and to the HC. Specifically, a strong correlation would indicate that the more RRO is derived by OB from rhythmic nasal airflow, the more it is transferred further, to its downstream targets. For the PFC, this was indeed the case; we found that dia-OB and OB-PFC coherences were positively correlated (Fig. 2C) in all sleep-wake states (R>0, p<0.1; Table S2), except SWS where the correlation was negative. In contrast, no such faithful, obligatory transmission of RRO through OB was found for the HC. No significant correlation between dia-OB and OB-HC coherences was detected in waking (see grey lines in Fig. 2C), and the correlation was negative in sleep states. Thus, the consistent increase in RRO conveyed by the OB to the PFC was closely associated with RRO variation in the OB network derived from respiration (Fig. 2C), whereas RRO transmission from OB to HC, varying in a wide range from one experiment to the next (Fig. 2B), did not follow the variations of dia-OB coherence in waking (AW and QW) and it was in fact showing an opposite tendency in sleep states (Fig. 2C).

### 3.3. Role of OB-mediated RRO in coupling between higher order networks in PFC and HC

Oscillatory coupling between PFC and HC was reported in different states, both in delta and theta frequency bands ^16, 20, 21^, as maintained by various mono- and polysynaptic connections between the two structures. Theta peaks were also dominant in PFC-HC coherence spectra during theta states (AW, REM sleep) in the present study. By adding simultaneous dia EMG and OB recordings, however, we also identify RRO coherence peaks indicative of PFC-HC coupling at RRO frequency in the delta range. They appeared either alone (QW) or in addition to theta (AW; Fig. 3A).

**Figure 3.**
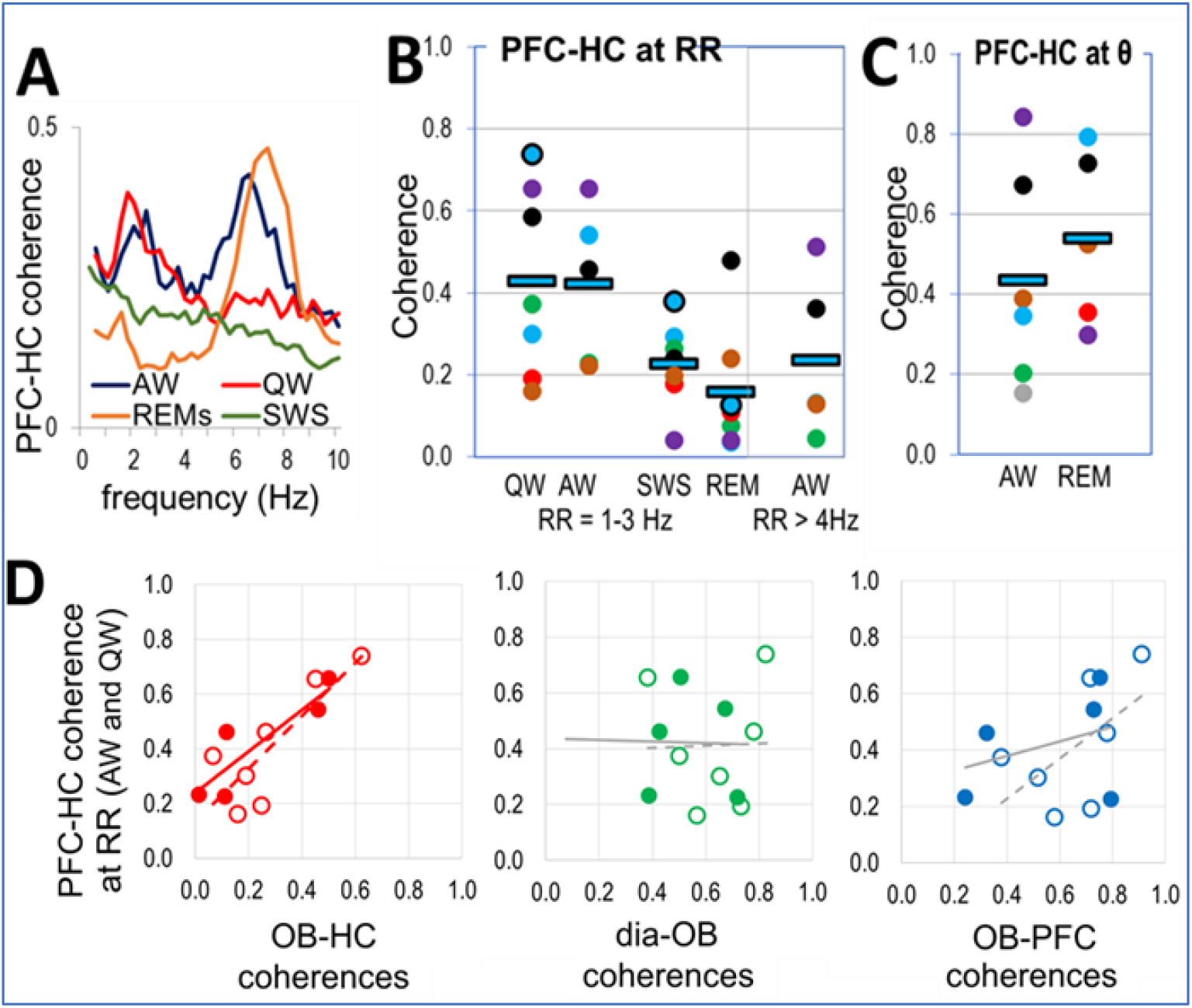
Coupling of PFC and HC networks by RRO. **A**. PFC-HC coherences in different states. **B-C**. PFC-HC coherences at RRO in awake (QW, AW) and sleep states (SWS, REM) at RRO (1-3 Hz) and at theta (6-8 Hz) frequencies in theta states (AW, REM). Squares: group averages, dots: individual experiments; same colors were used for individual rats in different states (same colors as in Fig. 2A). **D**. Correlation between peak RRO coherences connecting PFC and HC vs. RRO coherences connecting OB to HC, dia, and PFC signals in AW (filled symbols and solid trendlines) and QW (open symbols and dashed trendlines) recordings at RRO frequency RRO. Significant correlations are shown in the color of the corresponding dots, trendlines of non-significant correlations are shown in grey.

Average RRO coherence between PFC and HC fell in between OB-PFC and OB-HC coherence values in all states (Table 1C), although the differences were only significant in QW (p=0.02) – with a sufficient gap between OB-PFC and OB-HC coherences. PFC-HC coherence was significantly higher in waking (above 0.42) than sleep (below 0.22) and did not change significantly within these states (i.e. p>0.1 for AW vs. QW and REM vs. SWS comparisons) (Fig. 3B). The relationship of OB-HC < PFC-HC < OB-PFC coherences was robust; it was also valid in most individual experiments (60 and 71% of recordings in AW and QW, respectively). PFC-HC coherence in waking reached theta coherence in AW (0.44) and REM sleep (0.54) (Fig. 3C)

Unlike theta rhythm, primarily generated in HC and conveyed to PFC ^24^, producing strong PFC-HC coherence, the origin of the RRO coherence between these structures is less certain. RRO is generated outside of these structures, and from the OB it is faithfully transmitted to the PFC but much less reliably to the HC (Fig. 2A, 2C).

RRO coherence in the PFC-HC signal-pair may be due to RRO received by PFC from OB and then transmitted to HC or may be the effect of common input from OB to PFC and HC. The former appears consistent with relatively strong, state-dependent RRO in OB-PFC and PFC-HC coherences (compare Figs. 2A and 3B), and the latter with the large variability of RRO transmission to HC (Fig. 2B) in specific states, i.e. relatively high in some experiments and lower in others in wake states (Fig. 2A).

To distinguish between these possible mechanisms, we next compared correlations between RRO coherences in individual experiments in each state in which RRO coherences were present (Fig. 3D). We investigated in particular, whether stronger RRO synchrony between HC and PFC signals was associated with stronger OB-PFC or with stronger OB-HC coherences. As shown in Table 1B and Fig. 3D, PFC-HC RRO coherence significantly correlated only with RRO coherences linking OB with HC but not with PFC. This relationship was found in all states showing strong RRO. Thus, R^2^ correlation coefficient revealed similarity between the variations of PFC-HC and OB-HC coherences from one experiment to the next in QW (R^2^=0.69, p=0.003) and in AW (R^2^=0.79, p=0.01; Table 1B).

In contrast, PFC-HC coherence did not correlate with OB-PFC coherence in any state (Table 1B). A positive trend was observed in QW (R^2^=0.33) but was not significant (p=0.06; see grey lines in Fig. 3D). Variations of PFC-HC coherence in individual experiments was not affected by variations of dia-OB coherence in waking states and by any coherence connecting dia, OB, PFC, HC signals during sleep (Figs. 3D, S4).

## 4. Discussion

This study used inter-regional coherences and their correlations to trace the RRO signal from OB, where it is derived from rhythmic nasal airflow, to higher order brain networks of PFC and HC, where it may potentially contribute to communication between these structures by synchronizing their activities at the respiratory rate. We focused on on-going (i.e. “background”) RRO unaffected by behaviors requiring its short-term fluctuations, e.g. sniffing. We found that this rhythmicity depends on sleep-wake states; it is significantly larger in waking than in sleep. Within arousal states, however, it remains unchanged when the animal is engaged in behaviors or conditions, such as locomotion (AW), and REM sleep associated with theta vs. consummatory behaviors (QW) and SWS associated with non-theta HC activity. In agreement with previous reports ^3, 4, 9, 25^, RRO was more prominent in PFC with an obligatory transmission of RRO from OB to PFC. This was indicated by parallel variations in dia-OB and OB-PFC coherences in individual experiments, verified by significant correlation between these parameters over the group. In contrast, RRO was relatively low in HC, and the variations of OB-HC coherence did not necessarily follow those between dia and OB. RRO input to HC ^5-7^, however, lead to strong variations in individual recordings (at odds with the grossly defined states of AW and QW). Importantly, this variability, quantified with OB-HC coherence, was essential for establishing PFC-HC synchrony at the respiratory rate, whereas variations of RRO in OB and PFC had no significant effect.

### 4.1. RRO in waking

Communication and collaboration between HC and PFC and its impairment in psychiatric diseases has been the primary focus of extensive research to date ^18, 19, 24, 26^. HC-PFC theta synchronization and its role in spatial working memory is relatively well-studied ^27-32^. On the other hand, PFC is theorized to be the master regulator of working memory and higher-order executive function ^33-38^, yet the mechanisms by which PFC exerts “top-down” influences remain less clear. Functional coupling of PFC with downstream circuits, including HC, amygdala, ventral tegmental area, by means of delta-range oscillations (2-5 Hz) was recently shown in PFC-specific tasks ^14-16^. It was proposed that theta and delta oscillations may serve as parallel channels of communication between the HC and the PFC in opposite directions: theta HC-to-PFC and delta PFC-to-HC ^21^. The results of the present study indicate that the balance and interaction between theta and RRO may also provide a potential mechanism for bidirectional PFC-HC coupling. When RRO is within the delta range, it can contribute to or even drive the PFC 2-5 Hz rhythm ^39, 40^. PFC receives this input whenever OB is driven by RRO and may broadcast it widely. In contrast, the connection of HC to this global ^2^ rhythm appears more dynamic; when the HC is receptive to RRO input, PFC-to-HC channels would open in the delta range – distinct from the channel established by HC-theta in the opposite direction.

The most striking observation of this study was the marked contrast between the strength and reliability of RRO input, stronger in PFC than in HC (Table 1A), and its effect on PFC-HC coherence, i.e. RRO in HC more influential than in PFC (Table 1B). This raises questions regarding the mechanisms and the functional consequences. Differences in cytoarchitecture, connectivity, and other properties of PFC and HC networks give rise to different intrinsic oscillations in the two structures, which allow them to resonate with rhythmic input at specific frequencies. We propose that, due to these differences, baseline slow RRO recorded in this study may be involved in PFC-HC communication primarily in the of PFC-to-HC direction.

Task-related intrinsic oscillations in PFC appear at frequencies in the delta range ^14-16^, but these oscillations in waking are markedly different from the wide-band thalamo-cortical delta rhythm of SWS. Specifically, the delta (of waking) is spectrally of narrow-band, is hierarchically nested with gamma oscillations ^20, 41^, and is normally generated in cortico-cortical circuits, associated with various cognitive functions ^41-44^. Outgoing PFC messages may use the RRO fluctuations in sensitivity of downstream structures when PFC delta and RRO are synchronized. With this mechanism, RRO outside of sniffing ^5-7^ may make HC networks sensitive to messages arriving assembled in bouts at delta-range frequencies.

On the other hand, background RRO is slower than HC theta rhythm and thus, in order to synchronize with the signature HC oscillation during active states of sniffing, respiratory rate is accelerated and brought within the higher and narrower theta frequency band. The two oscillations, RRO and theta, show distinct characteristics in HC, such as different laminar profiles and theta-modulated gamma bands, and differentially entrain HC neurons ^5-7^ – even when their frequencies overlap. Olfactory-related activity patterns in OB, such as cell firing and gamma bursts, appear phase locked in these episodes to the synchronized RRO-theta rhythm ^45^. The exact mechanisms are not completely understood, but rhythmic synchronization of sensory sampling in OB on one hand and excitability of neurons involved in central processing in HC and piriform cortex on the other is considered a “paradigmatic example” of active sensing ^11, 46^. This would serve to optimize odor perception, coordinating it with multiple sensory channels, associated with rhythmic nasal, whisker, and head movements.

Although lacking strong direct projections from the OB ^6, 47, 48^, the PFC and HC receive RRO via multisynaptic pathways which includes common connections from the piriform cortex, as well as separate projections, through amygdala (PFC) or entorhinal cortex (HC)^3, 6^. Accordingly, the PFC-HC coherence may emerge from common OB input, or alternatively the RRO may be directed primarily to PFC and then transmitted to HC. Differential nodes in OB output pathways connecting PFC and HC (see e.g. ^21, 49^) may set the balance of RRO between the two structures.

When and how HC couples with slow baseline RRO will require further investigations using specific tasks beyond the sleep-wake states of this study. Data demonstrating the potential role of RRO in non-olfactory processing has accumulated in recent years, not only from rodent studies but also from human studies ^10, 50, 51^. A specific challenge for translating the results between species resides with important differences between human and rodents. Brain oscillations, functions, dynamics, and key features including characteristic frequencies are evolutionarily well-preserved ^52^, but respiratory rate varies widely between species. In humans, respiratory rate (∼0.2 Hz) is below the frequencies of the key components of the EEG oscillatory hierarchy which thus cannot establish coherent coupling with respiration. RRO remains however manifested in humans as respiratory modulation of the amplitude of brain oscillations including slow (delta, theta) as well as fast (beta, gamma) rhythms – involved in cognitive processes ^10^. This is a different form of coupling which unlike coherence does not require matching the frequencies of rhythms generated by different mechanisms.

### 4.2. RRO in sleep

In contrast to wake states, RRO during sleep appears reduced at the level of OB indicated by relatively low dia-OB coherence (Fig. 2) thus restricting OB-mediated RRO in higher brain structures (PFC, HC) from coupling with oscillations dominant in these networks during sleep. Viczko et al. ^53^ demonstrated for example that slow oscillations (SO), an archetypical EEG pattern in SWS, emerges separate from respiration even when they overlap in frequency, and argued that it “fits with an SO mechanism as intrinsic emergent property of a deafferented neural network”. Our data are consistent with this concept, suggesting that intrinsic brain oscillations – relevant in sleep-dependent memory consolidation both in SWS (SO ^53^ and delta ^54^) and REM sleep (theta ^55^) – are protected from RRO. It is interesting to note that in humans, where very few studies analyzed RRO during sleep with statistical scrutiny ^56, 57^, subtle changes in EEG linked to respiratory cycles were enhanced in SWS and REM sleep in children with sleep disordered breathing in multiple frequency bands, including delta ^56^, theta ^56, 57^, alpha, and sigma ^57^. Adenotonsillectomy, the most common surgical procedure for sleep apnea, which among other benefits improves cognitive function, reduced or normalized these RRO alterations.

It is important to note, however, that our conclusions only concern rhythmic RRO mediated by the OB. It has been reported that, besides RRO, respiration may also pace non-rhythmic events, linking their occurrence to specific phases of respiration (see e.g. ^13, 50^). In sleep, this may include sharp wave/ripples and dentate spikes; that is, intrinsic HC patterns during SWS synchronized with UP-DOWN transitions in cortical networks that are involved in functional PFC-HC interactions serving memory consolidation ^58, 59^. For instance, a recent study ^60^ found that the post-inspiratory bias of these patterns along with firing of a large number of PFC and HC neurons remained after deafferentation of OB sensory neurons, indicating that mechanisms that bypass the OB play a primary role in their synchronization. The authors hypothesized that the contribution of a “so-far undescribed ascending respiratory corollary discharge signal, likely propagating from the brainstem respiratory rhythm generators” could pace limbic networks using a disinhibition-mediated mechanism – consistent with lack of prominent LFPs in the absence of input from the OB ^60^. The causal model remains to be elucidated. In addition to ascending projections from the pre-Bötzinger complex ^61^ or the locus coeruleus ^62^, several other signals from internal organs may be involved –due to respiratory movements and chemosensitive signals from the cardiovascular system ^13, 63, 64^.

### 4.3. Potential relevance to COVID-19

We are not aware of published research on whether impaired RRO mechanisms are implicated in COVID-19 pathology. However, disturbances in smell, emerged early as a predominant neurological symptom ^65, 66^, serve as evidence for COVID-19 related neurological abnormalities originating from pathology of the olfactory epithelium. According to current understanding (rev. ^22, 67^), SARS-CoV-2 does not directly infect olfactory sensory neurons; their deficit is mediated instead by the altered microenvironment maintained by cells in the olfactory epithelium expressing ACE2 receptors ^68-70^. Yet, the potentially lasting damage is not localized to the OB; in the first prospective imaging studies (MRI scans 3-4 months after COVID-19 hospitalization), significant changes in grey matter volume were found, primarily in cingulate gyrus, piriform cortex, and hippocampus, correlated with loss of smell and also with memory loss^23^.

Much less is known about the cellular mechanism of RRO generation in the OB but conditions similar to those leading to smell loss may also affect RRO. Olfactory sensory neurons can respond not only to odorants but also to mechanical stimuli^71, 72^ and transmit both odor and air flow-driven mechanical signals^73, 74^. The latter was only studied however in mechanisms and schemes related to sniffing while their role in low frequency RRO, targeting a wide range of forebrain regions, remains unidentified. Involvement of non-sensory cells of the olfactory epithelium in RRO remains not clear either, but their impaired function in providing structural support, maintenance of ionic environments, etc. may negatively affect RRO, as well.

To date, we have solid data demonstrating that RRO depends on OB mechanisms ^2, 6, 8^ and modulates higher brain function ^10, 50, 51^. Abnormal PFC-HC coupling would not be immediately noticeable for patients, as the more obvious symptom of smell loss is, but it may lead to neurological consequences. Further studies are necessary to address the effect of SARS-CoV-2 on this system.

## Supplemental Information

**Figure S1.**
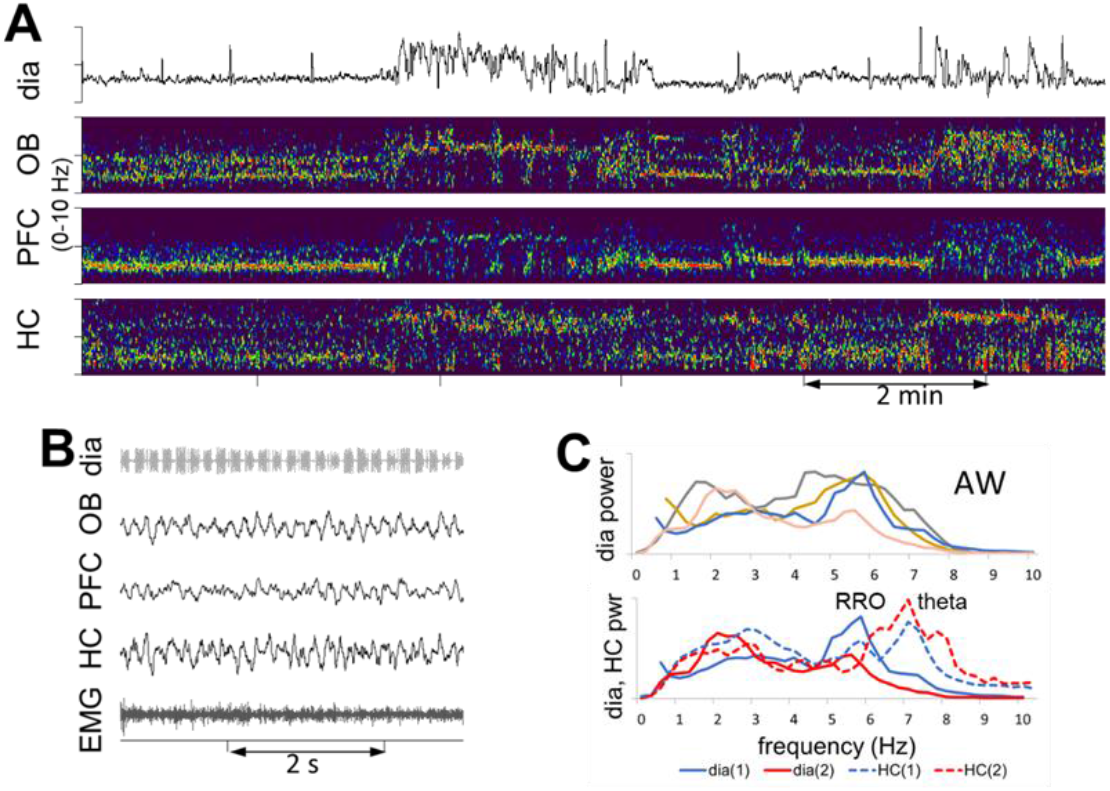
Unstable RRO in AW. Example of segments with RRO matching theta frequency, presumably associated with sniffing and excluded from analysis in this study (A) and segments with slow and faster RRO not synchronized with theta frequency (B). **A**. Rapidly alternating episodes of slow and fast respiration, presumably associated with sniffing. Respiration and time-frequency plots of LFP signals are shown in the 0-10 Hz range. Note matching frequencies of different signals in slow RRO and theta segments. **B**. Example of theta segment with RRO at theta frequency. **C**. *Top:* Two spectral peaks in dia autospectra, one at 2-3 Hz, the other at 4-6 Hz, in 4 out of 10 AW recordings. *Bottom:* Dia (solid lines) and HC (dashed lines) autospectra showing that high RRO did not overlap with HC theta.

**Figure S2.**
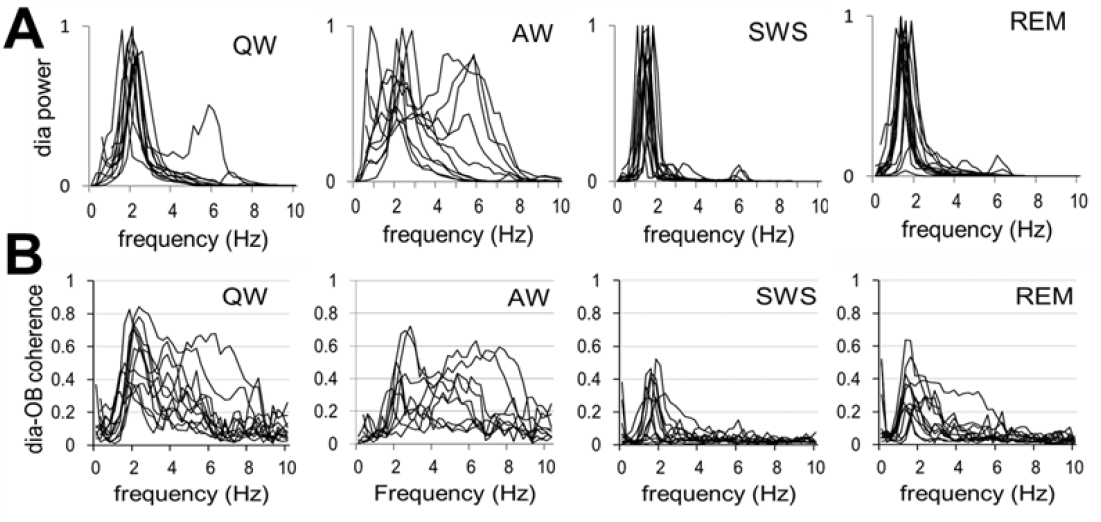
RRO oscillations in dia EMG, correlated with OB LFP. **A**. Autospectra of dia EMG in individual experiments. Note single sharp RRO peaks varying in a narrow frequency range in all states, except AW which showed larger variations. **B**. dia-OB coherence spectra in individual experiments. Additional peaks at higher frequencies appear also appear in dia-OB in a few recordings (1 in QW and 4 in AW) and more frequently in dia-OB coherences.

**Figure S3.**
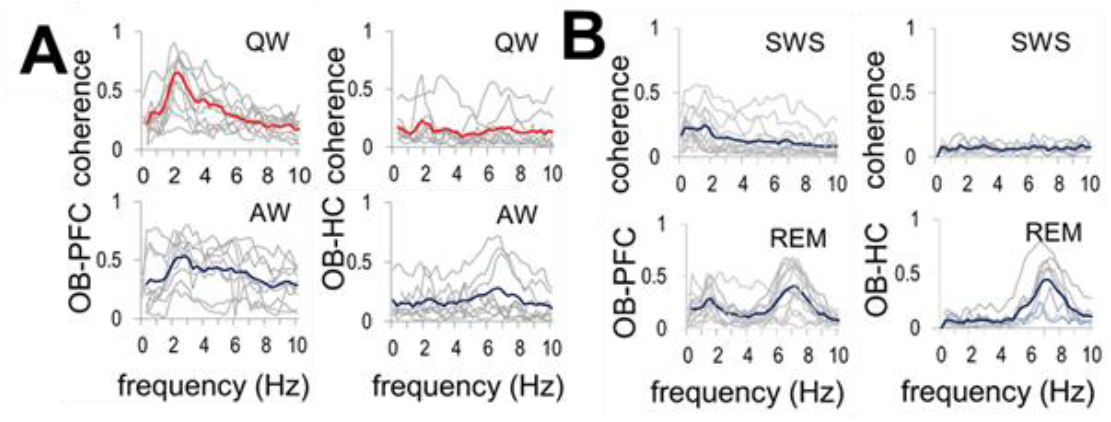
RRO coherence functions (0-10 Hz) between OB and higher order networks of PFC and HC during waking (**A**; QW and AW) and sleep (**B**; SWS and REM) in individual experiments (grey) and averaged over the group (red and blue). Note high RRO coherence in waking and low RRO coherence sleep, along with strong theta peak during REM sleep.

**Figure S4.**
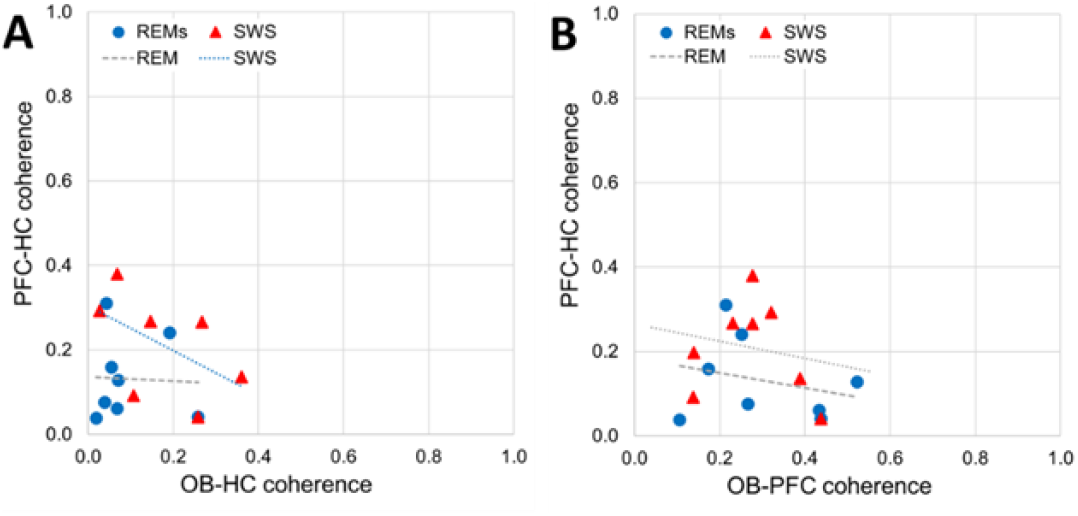
Correlation between peak RRO coherences between PFC and HC, and RRO coherences connecting OB to HC (**A**), and PFC signals (**B**) in different states with weak or non-existent RRO (REM sleep, SWS). Significant correlations are shown in the color of the corresponding dots, trendlines of non-significant correlations are shown in grey.

**Table S1:** length of segments and average RRO and theta frequency in different states

| Per rat (in 2 days): | SWS | REM | QW | AW |
| --- | --- | --- | --- | --- |
| # of segments | 9±0.8 | 9±0.9 | 7±0.4 | 7±0.5 |
| Ave length (s) | 127±11 | 109±12 | 112±18 | 95±14 |
| Total length (s) | 1237±219 | 1031±186 | 821±147 | 670±98 |
| # rats(recordings) with identified RRO | 7(14) | 7(14) | 7(12) | 6(9) |
| RRO peak freq. and range (Hz) | 1.65±0.05<br>1.2-2.0 | 1.65±0.05<br>1.2-2.0 | 2.01±0.08<br>1.5-2.4 | 2.23±0.11<br>1.0-2.9 |
| Theta freq. and range (Hz) |  | 7.08±0.09<br>6.8-7.3 |  | 6.84±0.13<br>6.6-7.1 |

**Table S2:** Correlation between dia-OB coherences and OB-PFC and OB-HC (p<0.1 positive correlations are marked with yellow background,). C(X-Y): coherence between X and Y.

| R2 between C(dia-OB) and... | SWS | REM | QW | AW |
| --- | --- | --- | --- | --- |
| ... C(OB-PFC) | 0.31 (-) | 0.17 | 0.33 | 0.21 |
| ... C(OB-HC) | 0.35 (-) | 0.41 (-) | 0.08 | 0.12 |

### Experiment procedures

Male rats (360–560g, Charles River Laboratories) were used in this study. Experiments were performed on 8 rats subjected to survival surgery followed by chronic recordings in free behaviors. All procedures were performed in accordance with the Institutional Animal Care and Use Committee of Beth Israel Deaconess Medical Center.

Survival surgeries were conducted in sterile conditions under deep anesthesia, maintained by a mixture of Ketamine-Xylazine (i/p; 30-40 mg/kg ketamine and 5 mg/kg Xylazine) with supplementary injections of Ketamine (10 % of the initial dose) if necessary. Stainless steel screws above the parietal cortex (AP: -3.5 mm, Lat: 2.5 mm from bregma) were inserted on the left side to record the cortical EEG and in the nasal bone (∼5.0 mm anterior to bregma) and above the cerebellum to act as ground and reference electrodes. Two single electrodes (stainless steel wires) were implanted on each side (AP: +3.2 mm, Lat: ±0.5 mm, DV: -5.1 mm) to record the electrical activity in the PFC, a single electrode on the left side for the OB (AP: +8.5 mm, Lat: 1.5 mm, DV: -1.6 mm), and a pair of twisted wires with 1 mm between their tips in the HC on the right side (AB: -3.7, Lat: 2.2 mm, DV: -3.5 mm) (Fig. S5). All the electrode wires and screws were fixed to the skull with dental acrylic. In addition, multi-threaded electrodes in soft insulation were implanted in the diaphragm to record diaphragmal EMG and in neck muscles to evaluate general motor activity and tone to identify sleep-wake states, and channeled under the skin to the head-connector. The electrodes were connected to an amplifier (A-M systems) for data recording, filtered between 0.3 and 100 Hz, whereas the dia EMG filters had both low- and high-pass filters set at 100 Hz to diminish unwanted noise including respiratory movement artifacts and electrical noise from the heart. Antibiotic gel was applied before suturing the incisions. Meloxicam analgesics, 1 mg/kg of 5 mg/ml daily, was given subcutaneously two days in a row to relieve pain. The rats were observed until they fully woke up. They recovered for a week in their home cages before the first 24 hours-recording. These recordings included different behaviors and sleep-wake states while the rat was freely moving in the cage, undisturbed. The rats were euthanized by 1 ml of Ketamine injections 3 weeks later and the brains were extracted for verification of the electrode position (Fig. S5).

### Data and statistical analysis

In two 24 hr recordings, acquired a couple of days apart in each rat, sleep-wake states were identified using standard criteria based on cortical EEG, HC LFP, and neck muscle EMG recordings. In waking, characterized by high and variable muscle activity, active waking (AW) and quiet waking (QW) were separated by the presence or absence, respectively, of HC theta rhythm and low amplitude fast cortical EEG. Sleep periods, characterized by continuously low muscle tone were also divided by the presence of HC theta accompanied by low-amplitude cortical EEG (REM sleep) or large amplitude slow activity dominating cortical and HC recordings (slow wave sleep, SWS). For analysis, multiple segments were selected from discontinuous episodes of each state dispersed over the 2 days of recordings, in which respiration appeared relatively stable without fluctuations (Table S1). Respiratory rate varied in different states in a relatively narrow band (between 1-3 Hz, Table S1) with an occasional faster component (4-6 Hz) in AW which did not overlap with theta frequency, specifically verified in each segment submitted for analysis.

EEG (filtered between 0.3 and 100 Hz) and EMG (100-300 Hz) recordings were acquired as a ∼.DDF-files in DASYLab 7.0 (at 1 kHz sampling rate) and then imported to Spike2 (Version 7.06, Cambridge Electronic Devices) for signal analysis. Dia EMG recordings were processed using built-in procedures of Spike2 to remove ECG contamination and to convert high-frequency EMG components to retrieve pure respiratory rhythm.

Noise-free segments with stable respiration for at least ∼100 s were selected in SWS, REM, QW and AW, recorded on two different days (see Table S1 for the number and length of segments in each state), and submitted to Fast Fourier Transform (FFT) to obtain power spectra and coherence functions. FFT was performed on consecutive 4.096 s windows, using a program (COHER.S2S) from the Spike2 library. FFT converts a block of waveform data into frequency representation; the FFT of a block of n data points sampled at a frequency F produces n/2+1 equally spaced complex numbers (A_f_ = a e^jφ^), from a frequency of 0 to F/2, spaced by F/n. Power spectral density of each signal and cross spectral density for signal pairs are calculated using these complex numbers, as psd(f) = a e^jφ^ a e^-jφ^ = a^2^ and csd(f) = a e^jφ(a)^ b e^-jφ(b)^ = a b e^j(φa-φb)^. To quantify neuronal synchronization we used coherence analysis calculated using these psd and csd-s, as coh(f) = (SUM(csd_ab_(f))^2^ / SUM(psd_a_(f)) * SUM(psd_b_(f)), where SUM(.) denotes summation over blocks of data.

Coherence function is calculated for pairs of different signals of interest (e.g., LFP recorded from two spatially distinct areas) and provides a useful measure of synchrony between two signals at each frequency of the spectrum, thus revealing characteristic frequencies of synchronization. Pairwise coherences were calculated between 5 signal pairs, representing the potential transfer of the RRO signal to higher-order structures through the OB (i.e. dia with OB and OB with PFC and HC) and between these higher order structures (i.e. PFC with HC). Power spectra for dia EMG and HC were also calculated to identify the frequencies of spectral peaks of RRO and theta rhythm. Coherence values were compared against chance using surrogate-based statistical testing by computing coherence spectra between PFC and HC vs. OB of the same rat as well as OB from a different animal recorded at the same time and processed in parallel using different channels of the same amplifier, A/D converter, and stored in the same file. The significance of coherence values depends on the length of the analyzed segment (see different levels of erroneous coherence in examples of short and long segments in Fig. S6A-B) but was statistically different (lower) in all signal pairs (p<0.0001 for OB-PFC and PFC-HC in all states, as well as for OB-HC in QW and SWS; p=0.025 and p=0.001 for OB-HC in REM and AW, respectively). To verify that the RRO coherences were not driven by the common reference over the cerebellum, cross-correlation was calculated on a QW segment which showed high RRO coherence, in each rat. The phases between different LFPs were inconsistent between experiments, but the RRO peaks did not overlap (see examples in Fig. S6C-E). Differences between coherences in different states were tested using Student’s t-test after Fisher r to z transformation to obtain z-scored values with normal distribution. Correlation between pair-wise coherences were statistically tested using Excel’s T-DIST procedure using the formula p=TDIST(R*SQRT((N-2)/(1-R2)),N,1) where R is coherence and N is number of experiments, included.

### Perfusion and Histology

Perfusion was conducted by inserting a hypodermic needle connected to a pump into the ascending aorta through the left ventricle of the heart. The right ventricle was cut open and phosphate buffered saline (PBS) was pumped into the circulation system for 5 minutes, until all blood had been drained and replaced with PBS. Then 10% buffered formalin solution was pumped for about 15-20 minutes, until the rat was stiff. After perfusion, the rats’ brain was carefully extracted and placed in a vial filled with 20 mL of formalin for further fixation for at least one hour, while stored at 4°C. Before cryosectioning (Microm, HM 450), the brains were stored in 20% sucrose solution overnight at 4 °C until sank to the bottom of the vial. 45 µm slices were stored in PBS with Azide and then mounted on slides and stained with Cresyl Violet, coverslipped and evaluated to locate the marks of damage due to inserted recording electrodes (Fig. S5).

**Figure S5.**
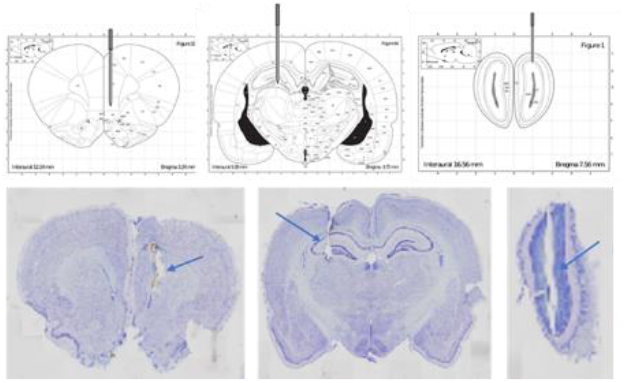
Placement of electrodes for local field potential (LFP) recordings. Schematic drawing (*top*) and histology of stained coronal sections (*bottom*) to identify electrode coordinates in the prefrontal cortex (PFC), hippocampus (HC), and olfactory bulb (OB).

**Figure S6.**
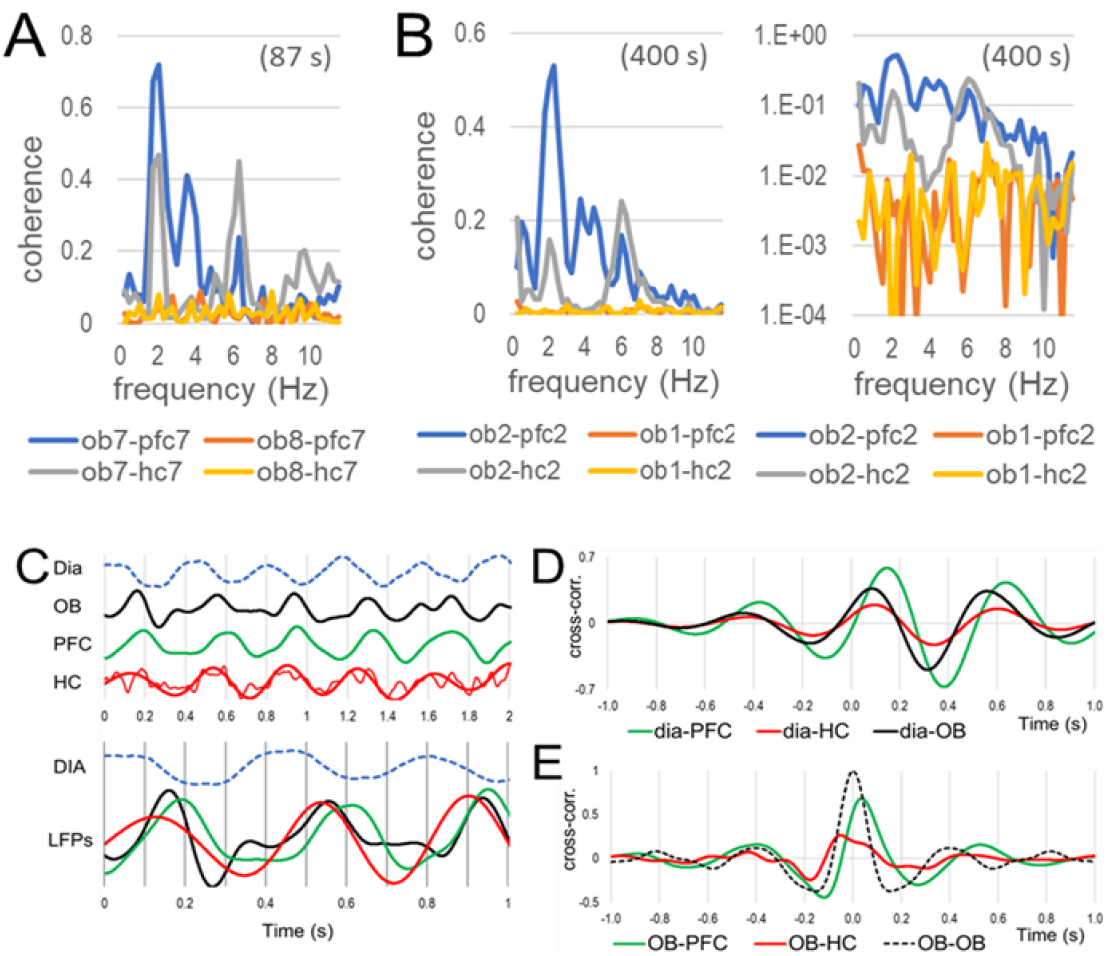
Coherence analysis. **A-B**. Comparison of OB-PFC (*blue*) and OB-HC (*grey*) coherences with coherences values calculated using a surrogate signal (OB of a different rat, recorded simultaneously, *red* and *yellow*) in a short (A) and long segment (B, shown in linear and log scale). **C-E**. Phase relationship between RRO in different LFPs. **C**. *Top*: Short (2 s) sample recording of dia, OB, PFC, and HC in QW. LFPs were filtered below 18 Hz (HC was additionally filtered below 3 Hz to obtain a visually smooth trace for illustration). *Bottom*: same traces (LFPs overlapped) on different time scale. **D**. Sample cross-correlograms between dia EMG and PFC, HC, and OB LFPs in QW. **E**. OB autocorrelogram (dashed line) and cross-correlogram between OB and higher order (PFC and HC) LFPs. B and C are from the same QW segment.

## Acknowledgements

This work was supported by the National Institute of Mental Health grant R01 MH100820 to BK and a Swedish Government stipend to RM.

## Author contributions

RM and BK contributed to the experiments, analysis, interpreting the results, and writing the report.

## References

1. Adrian, E.D. Olfactory reactions in the brain of the hedgehog. J Physiol 100, 459–473 (1942).

2. Tort, A.B.L., Brankack, J. & Draguhn, A. Respiration-Entrained Brain Rhythms Are Global but Often Overlooked. Trends Neurosci 41, 186–197 (2018).

3. Biskamp, J., Bartos, M. & Sauer, J.F. Organization of prefrontal network activity by respiration-related oscillations. Sci Rep 7, 45508 (2017).

4. Zhong, W., et al. Selective entrainment of gamma subbands by different slow network oscillations. Proc Natl Acad Sci U S A 114, 4519–4524 (2017).

5. Nguyen Chi, V., et al. Hippocampal Respiration-Driven Rhythm Distinct from Theta Oscillations in Awake Mice. J Neurosci 36, 162–177 (2016).

6. Yanovsky, Y., Ciatipis, M., Draguhn, A., Tort, A.B. & Brankack, J. Slow oscillations in the mouse hippocampus entrained by nasal respiration. J Neurosci 34, 5949–5964 (2014).

7. Lockmann, A.L., Laplagne, D.A., Leao, R.N. & Tort, A.B. A Respiration-Coupled Rhythm in the Rat Hippocampus Independent of Theta and Slow Oscillations. J Neurosci 36, 5338–5352 (2016).

8. Ito, J., et al. Whisker barrel cortex delta oscillations and gamma power in the awake mouse are linked to respiration. Nat Commun 5, 3572 (2014).

9. Tort, A.B.L., et al. Parallel detection of theta and respiration-coupled oscillations throughout the mouse brain. Sci Rep 8, 6432 (2018).

10. Zelano, C., et al. Nasal Respiration Entrains Human Limbic Oscillations and Modulates Cognitive Function. J Neurosci 36, 12448–12467 (2016).

11. Corcoran, A.W., Pezzulo, G. & Hohwy, J. Commentary: Respiration-Entrained Brain Rhythms Are Global but Often Overlooked. Front Syst Neurosci 12, 25 (2018).

12. Heck, D.H., Kozma, R. & Kay, L.M. The rhythm of memory: how breathing shapes memory function. J Neurophysiol 122, 563–571 (2019).

13. Heck, D.H., et al. Breathing as a Fundamental Rhythm of Brain Function. Front Neural Circuits 10, 115 (2016).

14. Dejean, C., et al. Prefrontal neuronal assemblies temporally control fear behaviour. Nature 535, 420–424 (2016).

15. Karalis, N., et al. 4-Hz oscillations synchronize prefrontal-amygdala circuits during fear behavior. Nat Neurosci 19, 605–612 (2016).

16. Fujisawa, S. & Buzsaki, G. A 4 Hz oscillation adaptively synchronizes prefrontal, VTA, and hippocampal activities. Neuron 72, 153–165 (2011).

17. Buzsaki, G. Theta oscillations in the hippocampus. Neuron 33, 325–340 (2002).

18. Sigurdsson, T., Stark, K.L., Karayiorgou, M., Gogos, J.A. & Gordon, J.A. Impaired hippocampal-prefrontal synchrony in a genetic mouse model of schizophrenia. Nature 464, 763–767 (2010).

19. Dickerson, D.D., Wolff, A.R. & Bilkey, D.K. Abnormal long-range neural synchrony in a maternal immune activation animal model of schizophrenia. J Neurosci 30, 12424–12431 (2010).

20. Pittman-Polletta, B., Hu, K. & Kocsis, B. Subunit-specific NMDAR antagonism dissociates schizophrenia subtype-relevant oscillopathies associated with frontal hypofunction and hippocampal hyperfunction. Sci Rep 8, 11588 (2018).

21. Roy, A., Svensson, F.P., Mazeh, A. & Kocsis, B. Prefrontal-hippocampal coupling by theta rhythm and by 2-5 Hz oscillation in the delta band: The role of the nucleus reuniens of the thalamus. Brain Struct Funct 222, 2819–2830 (2017).

22. Cooper, K.W., et al. COVID-19 and the Chemical Senses: Supporting Players Take Center Stage. Neuron 107, 219–233 (2020).

23. Lu, Y., et al. Cerebral Micro-Structural Changes in COVID-19 Patients - An MRI-based 3-month Follow-up Study. EClinicalMedicine 25, 100484 (2020).

24. Nandi, B., Swiatek, P., Kocsis, B. & Ding, M. Inferring the direction of rhythmic neural transmission via inter-regional phase-amplitude coupling (ir-PAC). Sci Rep 9, 6933 (2019).

25. Heck, D.H., et al. Cortical rhythms are modulated by respiration. BioRxiv, 049007 (2016).

26. Lee, H., et al. Early cognitive experience prevents adult deficits in a neurodevelopmental schizophrenia model. Neuron 75, 714–724 (2012).

27. Sarnthein, J., Petsche, H., Rappelsberger, P., Shaw, G.L. & von Stein, A. Synchronization between prefrontal and posterior association cortex during human working memory. Proc Natl Acad Sci U S A 95, 7092–7096 (1998).

28. Igarashi, K.M. Plasticity in oscillatory coupling between hippocampus and cortex. Curr Opin Neurobiol 35, 163–168 (2015).

29. Zhang, X., et al. Impaired theta-gamma coupling in APP-deficient mice. Sci Rep 6, 21948 (2016).

30. Kaplan, R., et al. Medial Prefrontal-Medial Temporal Theta Phase Coupling in Dynamic Spatial Imagery. J Cogn Neurosci 29, 507–519 (2017).

31. Tamura, M., Spellman, T.J., Rosen, A.M., Gogos, J.A. & Gordon, J.A. Hippocampal-prefrontal theta-gamma coupling during performance of a spatial working memory task. Nat Commun 8, 2182 (2017).

32. Barr, M.S., et al. Impaired theta-gamma coupling during working memory performance in schizophrenia. Schizophr Res 189, 104–110 (2017).

33. Teffer, K. & Semendeferi, K. Human prefrontal cortex: evolution, development, and pathology. Prog Brain Res 195, 191–218 (2012).

34. Fletcher, P.C., Shallice, T., Frith, C.D., Frackowiak, R.S. & Dolan, R.J. The functional roles of prefrontal cortex in episodic memory. II. Retrieval. Brain 121 (Pt 7), 1249–1256 (1998).

35. Fletcher, P.C., Shallice, T. & Dolan, R.J. The functional roles of prefrontal cortex in episodic memory. I. Encoding. Brain 121 (Pt 7), 1239–1248 (1998).

36. Buckner, R.L. & Wheeler, M.E. The cognitive neuroscience of remembering. Nat Rev Neurosci 2, 624–634 (2001).

37. Miller, E.K. & Cohen, J.D. An integrative theory of prefrontal cortex function. Annu Rev Neurosci 24, 167–202 (2001).

38. Benchenane, K., Tiesinga, P.H. & Battaglia, F.P. Oscillations in the prefrontal cortex: a gateway to memory and attention. Curr Opin Neurobiol 21, 475–485 (2011).

39. Kocsis, B., Pittman-Polletta, B.R. & Roy, A. Respiration-coupled rhythms in prefrontal cortex: beyond if, to when, how, and why. Brain Struct Funct 223, 11–16 (2018).

40. Lockmann, A.L.V. & Tort, A.B.L. Nasal respiration entrains delta-frequency oscillations in the prefrontal cortex and hippocampus of rodents. Brain Struct Funct 223, 1–3 (2018).

41. Hunt, M.J., Kopell, N.J., Traub, R.D. & Whittington, M.A. Aberrant Network Activity in Schizophrenia. Trends Neurosci 40, 371–382 (2017).

42. Nacher, V., Ledberg, A., Deco, G. & Romo, R. Coherent delta-band oscillations between cortical areas correlate with decision making. Proc Natl Acad Sci U S A 110, 15085–15090 (2013).

43. Riecke, L., Sack, A.T. & Schroeder, C.E. Endogenous Delta/Theta Sound-Brain Phase Entrainment Accelerates the Buildup of Auditory Streaming. Curr Biol 25, 3196–3201 (2015).

44. Hall, T.M., de Carvalho, F. & Jackson, A. A common structure underlies low-frequency cortical dynamics in movement, sleep, and sedation. Neuron 83, 1185–1199 (2014).

45. Kay, L.M., et al. Olfactory oscillations: the what, how and what for. Trends Neurosci 32, 207–214 (2009).

46. Wachowiak, M. All in a sniff: olfaction as a model for active sensing. Neuron 71, 962–973 (2011).

47. Hoover, W.B. & Vertes, R.P. Anatomical analysis of afferent projections to the medial prefrontal cortex in the rat. Brain Struct Funct 212, 149–179 (2007).

48. Moberly, A.H., et al. Olfactory inputs modulate respiration-related rhythmic activity in the prefrontal cortex and freezing behavior. Nat Commun 9, 1528 (2018).

49. Bagur, S. & Benchenane, K. Taming the oscillatory zoo in the hippocampus and neo-cortex: a review of the commentary of Lockmann and Tort on Roy et al. Brain Struct Funct 223, 5–9 (2018).

50. Perl, O., et al. Human non-olfactory cognition phase-locked with inhalation. Nat Hum Behav 3, 501–512 (2019).

51. Arshamian, A., Iravani, B., Majid, A. & Lundstrom, J.N. Respiration Modulates Olfactory Memory Consolidation in Humans. J Neurosci 38, 10286–10294 (2018).

52. Buzsaki, G. & Draguhn, A. Neuronal oscillations in cortical networks. Science 304, 1926–1929 (2004).

53. Viczko, J., Sharma, A.V., Pagliardini, S., Wolansky, T. & Dickson, C.T. Lack of respiratory coupling with neocortical and hippocampal slow oscillations. J Neurosci 34, 3937–3946 (2014).

54. Kim, J., Gulati, T. & Ganguly, K. Competing Roles of Slow Oscillations and Delta Waves in Memory Consolidation versus Forgetting. Cell 179, 514–526 e513 (2019).

55. Boyce, R., Glasgow, S.D., Williams, S. & Adamantidis, A. Causal evidence for the role of REM sleep theta rhythm in contextual memory consolidation. Science 352, 812–816 (2016).

56. Chervin, R.D., et al. Correlates of respiratory cycle-related EEG changes in children with sleep-disordered breathing. Sleep 27, 116–121 (2004).

57. Immanuel, S.A., et al. Respiratory cycle-related electroencephalographic changes during sleep in healthy children and in children with sleep disordered breathing. Sleep 37, 1353–1361 (2014).

58. Sirota, A. & Buzsaki, G. Interaction between neocortical and hippocampal networks via slow oscillations. Thalamus Relat Syst 3, 245–259 (2005).

59. Sirota, A., Csicsvari, J., Buhl, D. & Buzsaki, G. Communication between neocortex and hippocampus during sleep in rodents. Proc Natl Acad Sci U S A 100, 2065–2069 (2003).

60. Karalis, N. & Sirota, A.M. Breathing coordinates limbic network dynamics underlying memory consolidation. BioRxiv (2018).

61. Yang, C.F. & Feldman, J.L. Efferent projections of excitatory and inhibitory preBotzinger Complex neurons. J Comp Neurol 526, 1389–1402 (2018).

62. Yackle, K., et al. Breathing control center neurons that promote arousal in mice. Science 355, 1411–1415 (2017).

63. Kocsis, B., Karlsson, T. & Wallin, B.G. Cardiac- and noncardiac-related coherence between sympathetic drives to muscles of different human limbs. Am J Physiol 276, R1608–1616 (1999).

64. Gebber, G.L., Barman, S.M. & Kocsis, B. Coherence of medullary unit activity and sympathetic nerve discharge. Am J Physiol 259, R561–571 (1990).

65. Boscolo-Rizzo, P., et al. Evolution of Altered Sense of Smell or Taste in Patients With Mildly Symptomatic COVID-19. JAMA Otolaryngol Head Neck Surg (2020).

66. Menni, C., et al. Real-time tracking of self-reported symptoms to predict potential COVID-19. Nat Med 26, 1037–1040 (2020).

67. Iadecola, C., Anrather, J. & Kamel, H. Effects of COVID-19 on the Nervous System. Cell 183, 16–27 e11 (2020).

68. Ziegler, C.G.K., et al. SARS-CoV-2 Receptor ACE2 Is an Interferon-Stimulated Gene in Human Airway Epithelial Cells and Is Detected in Specific Cell Subsets across Tissues. Cell 181, 1016–1035 e1019 (2020).

69. Li, W., Li, M. & Ou, G. COVID-19, cilia, and smell. FEBS J (2020).

70. Rockx, B., et al. Comparative pathogenesis of COVID-19, MERS, and SARS in a nonhuman primate model. Science 368, 1012–1015 (2020).

71. Grosmaitre, X., Santarelli, L.C., Tan, J., Luo, M. & Ma, M. Dual functions of mammalian olfactory sensory neurons as odor detectors and mechanical sensors. Nat Neurosci 10, 348–354 (2007).

72. Connelly, T., et al. G protein-coupled odorant receptors underlie mechanosensitivity in mammalian olfactory sensory neurons. Proc Natl Acad Sci U S A 112, 590–595 (2015).

73. Iwata, R., Kiyonari, H. & Imai, T. Mechanosensory-Based Phase Coding of Odor Identity in the Olfactory Bulb. Neuron 96, 1139–1152 e1137 (2017).

74. Carey, R.M., Verhagen, J.V., Wesson, D.W., Pirez, N. & Wachowiak, M. Temporal structure of receptor neuron input to the olfactory bulb imaged in behaving rats. J Neurophysiol 101, 1073–1088 (2009).

